# Combinatorial factorizable libraries outperform enumerated and random libraries for antibody discovery

**DOI:** 10.1101/2025.09.25.678628

**Authors:** Zheng Dai, Jonathan Krog, Emeline Labatut, Christine Banholzer, Michael Birnbaum, Stefan Ewert, David K Gifford

## Abstract

The effective use of high-throughput functional assays to discover novel biological therapeutics requires a diverse library of desirable candidates. Here we design candidate antibody libraries using factorizable neural networks (FNNs) that permit the economic synthesis of factorizable libraries that are depleted of non-specific binders. We experimentally evaluated an FNN designed antibody CDR-H3 factorizable library that is designed to contain binders to peptide-MHC (pMHC) complexes with depletion of library sequences that exhibit non-specific pMHC binding. The factorizable library comprises 10^4^ prefixes that were randomly combined with 10^4^ suffixes. We find that it contains more viable candidates than designed libraries with the same DNA synthesis budget on our pMHC binding task. FNNs make it possible to efficiently computationally design combinatorial libraries, a task that is not possible with conventional methods.

## 1 Introduction

High-throughput display platforms are increasingly being used to discover biologics such as antibody therapeutics. These display platforms express diverse candidate libraries using phage or yeast display followed by multiple rounds of affinity selection against a biological target of interest [11]. Therapeutic discovery by selection requires the design of libraries with candidates that possess desirable properties, such as affinity towards a given class of interest, while avoiding undesirable ones, such as polyspecificity, hydrophobicity, and instability [12]. The design of ideal candidate libraries has been an area of recent interest as a result of their practical importance as well as the computational challenges involved in their design [10, 13, 4].

Libraries of candidate antibodies typically contain randomly generated candidates or individually designed candidates. Most contemporary antibody candidate libraries are *random libraries* that are produced via mutagensis or sequential synthesis using trinucleotide codon mixes. While highly diverse, random libraries can be inefficient and contain sequences with undesirable qualities. An alternative to random libraries are rationally designed libraries where each library sequence is individually specified and synthesized, which we refer to here as *enumerated libraries*. Sequences in enumerated libraries can have superior developability profiles, resulting in efficient therapeutic discovery [10, 13]. However, enumerated libraries can be costly to manufacture, and the cost of enumerated libraries is prohibitive for the complexities we explore in this work. Recent results have used structural models to design antibodies to bind specific targets, but these approaches depended upon the knowing accurate target structures and still require enumerated library based screens and randomized optimization for final candidate selection and have not been utilized to create libraries of binders against a target class [6, 2]. The approach described here does not depend upon pre-existing knowledge of target structures or target structural prediction.

In previous work, we proposed the combinatorial composition of enumerated libraries, called *factorizable libraries* [4]. In this approach, relatively small libraries of prefix and suffix segments are rationally designed and then randomly ligated together to form an exponentially larger factorizable library, achieving both the benefits of having higher concentrations of desirable candidates via rational design and of high library complexity via random ligation. We focus our attention on the design of the hypervariable *complementarity-determining region 3* of the antibody heavy chain (CDR-H3) as it plays a major role in determining antibody binding and specificity [17]. We note the formation of variable CDR-H3 sequences *in vivo* also starts with the random joining of a set of germline encoded segments [15].

We find in this work that factorizable libraries produce a superior number of specific peptide-MHC binding antibodies when compared with random and enumerated library controls with the same DNA synthesis budget. In our experimental investigation desired library candidates are monoclonal antibodies (mAbs) with variable CDR-H3 regions that are depleted of non-specific peptide-MHC complex (pMHC) binders. We introduce a practical method of designing factoriable libraries using what we call a *factorizable neural network*, which allows us to efficiently and exactly compute the expected value of an objective function on the random combination of a set of candidate prefixes and a set of candidate suffixes. Finally, to create objective functions for factorizable neural networks we integrate heterogenous phage display data to learn antibody-target affinities and quantify the uncertainty associated with the affinities, which can then be used to guide library design. Importantly, this allows us to quantify and design libraries against non-specificity.

## 2 Methods

### 2.1 Computation of specific and non-specific affinities of CDR-H3 sequences from display platforms

We use data from phage display experiments to infer antibody-target affinities for each CDR-H3 sequence in a library. Phage display allow the assessment of affinities of up to 10^10^ unique antibodies in a single experiment [1]. Phage display works as follows: a library of biologics, in our case monoclonal antibodies (mAbs) are cloned into a phagemid vector template and expressed in bacteriophages. The phages display the mAbs on their surface, and they are then exposed to a target of interest. Antibodies that bind to the target (for example a pMHC) can be purified away from non-binding phages. The result of this selection is used as the input for the next selection, and these *rounds* are iterated until the remaining phages display antibodies with high affinity to the target. At the conclusion of each selection round the resulting antibody library of phages is sequenced. Sequencing reads are mapped to the vector template to deduce the identities and counts of the CDR-H3 sequences after each round of selection (Methods 5.2).

Different targets can be selected against in different rounds, which can be used to obtain polyspecificity data. For example, if we select mAbs against target A (Round 1), target A (Round 2), and target B (Round 3) we will select for mAbs sequences that bind both target A and target B. Alternatively, while selecting against a given target a second unimmobilized target can be added to the round to compete against the primary target. The second target will be washed away at the end of the round and carry mAbs with high affinity towards it with it. We refer to this as *counterselection*, and it provides a secondary method of assessing affinity.

We extract antibody affinity values from selection-based read counts (Methods 5.3). Read counts of CDR-H3 sequences after selection provide critical information about antibody properties. Previous studies have used simple heuristics to determine which antibodies are promising, such as a round-over-round CDR-H3 read count increase or the presence of CDR-H3 sequences after a final selection step. We compute the likelihood of CDR-H3 read counts given the affinities of each antibody to each target based on a model of binding kinetics. We apply variational inference to solve the inverse problem, yielding antibody affinities (Methods 5.3).

We further decompose mAb affinity into specific and non-specific components by simultaneously solving for these components in the context of multiple experiments that have been designed to resolve specificity using varied targets in each selection round (Methods 5.4). A key advantage of our model based approach to attaining affinity values is that it allows us to take phage display data with heterogeneous targets and counterselection targets and infer affinity using a single consistent model. This integration of heterogeneous target affinity information further allows us to decompose our affinity values into specific and non-specific components at the inference stage.

### 2.2 Objective function guided design of factorizable libraries using Factorizable Neural Networks

We design a diverse *factorizable library* [4] of mAbs using a novel objective that depletes non-specific pMHC binders to test the hypothesis that this library will produce more specific pMHC binders than random or enumerated libaries with identical DNA synthesis budgets. In a factorizable library, each sequence is a combination of independent *segments* from designed *segment libraries*. We use two segment libraries: a prefix segment library and a suffix segment library. The random combination of prefix and suffix segments allows the library size to scale up quadratically compared to the sizes of the segment libraries. Once segment libraries of DNA oligonucleotides are designed and synthesized, they can be joined using blunt end ligation, which generates the final factorizable library [3, 5].

Prior work has shown that the computational design of segment libraries of size *n* that are randomly combined is a computationally intractable problem. This is because a naive solution computes an objective function by evaluating all *n*^2^ factorizable library members. Thus, recomputation of the overall objective function is required when optimizing a single prefix or suffix candidate sequence. Ideally we would like to be able to quickly recompute our objective function when updating a single sequence with a simple update.

To enable efficient objective function updates and solve our combinatorial impasse we adopt an approach called the *reverse kernel trick* [4]. In this approach, we design our optimization objective so that the objective value for a sequence *f* (*s*), in our case the CDR-H3 sequence of a mAb, is expressed as an inner product of a pair of embeddings:

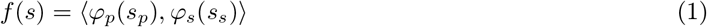

where *s* is the concatenation of *s*_*p*_ and *s*_*s*_, and *φ*_*p*_ and *φ*_*s*_ are prefix embedding functions that map prefix and suffix segments to some inner product space. Note that we consider the sequences and the segments to be of fixed length, and we model variable sequence length through padding. We hold the mAb framework constant in all domains except CDR-H3, and thus CDR-H3 affinities can be represented as a function that inputs a string of amino acids representing the CDR-H3 domain and outputs a real value. We can compute the objective over an entire factorizable library:

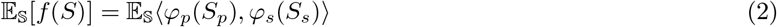

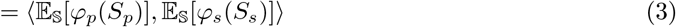

where *S* ranges over the sequences in the factorizable library. The factorization in the second step is consequence of the independence of segment assortment and the linear properties of the inner product. While any inner product space and inner product can be used, in this work we use ℝ ^*n*^ equipped with the dot product as our space.

The use of an inner product between prefix and suffix embeddings allows us to evaluate the average value of an objective without embedding all possible combinations of prefixes with suffixes, which is quadratic (𝒪 (*n*^2^)) in the number of prefix and suffix segments. Instead, we only need to embed the prefix library and the suffix library, which we can do in linear time (𝒪 (*n*)). Furthermore, if we store the sequence embeddings we can evaluate the result of changing one segment in constant time (𝒪 (1)) by only reembedding that single sequence.

We will represent functions of CDR-H3 affinity using a *Factorizable Neural Network* (FNN) that can be expressed as an inner product of prefix and suffix embeddings. FNNs are neural networks with the following functional form:

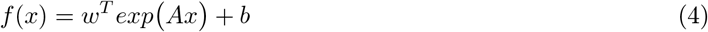

where *x* is the input, *A* is the matrix of weights between the input and a hidden layer with *h* units, *w* is the vector of weights between the hidden and output layer, and *b* is some scalar bias. *exp*(.) is used to denote the entrywise exponential function. We note that this is a three layer perceptron with exponential activation.

The key feature of this architecture is that if an input signal is made up of a linear combination of different vectors, then this architecture can factorize its output. Suppose *x* = ∑_*i*_ *x*_*i*_. Then:

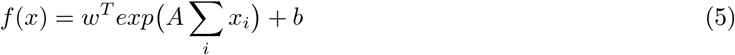

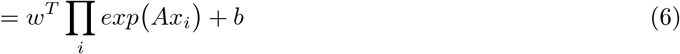

where ∏ _*i*_ *exp*(*Ax*_*i*_) is used to denote an entrywise product.

A factorizable neural network permits us to factorize *x* when it is a concatenation of one-hot vectors that encode a sequence. To construct this encoding, we first fix a map between an alphabet *Σ* and one-hot vectors. One-hot vectors are elements of ℝ ^|*Σ*|^ where exactly one entry is equal to 1 and all other entries are equal to 0. To encode a sequence of length *L* as a vector, we first map each character in the sequence to its one-hot vector. We then concatenate the vector, attaining a representation of length |*Σ*| *L*. To represent a prefix, we map each character to one-hot vectors, concatenate them, and the pad out the remainder of the vector with 0s until we get to |*Σ*| *L*. To represent a suffix, we map each character to one-hot vectors, concatenate them, and prefix the string with enough 0s to get to |*Σ*| *L*. Under this representation, the concatenation of the prefix and suffix in sequence space is simply the sum of the prefix and suffix in the encoding space.

This factorization allows us to immediately derive prefix and suffix feature maps, where the dimension of feature space of the embedding functions is the number of units in the hidden layer of the FNN. Let *x*_*p*_ and *x*_*s*_ denote the prefix and suffix encoding of *x* respectively. Then:

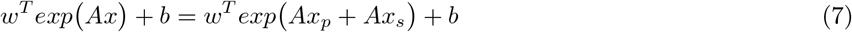

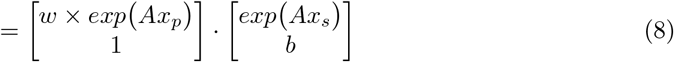

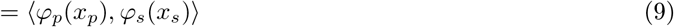

where “× “ here denotes entrywise multiplication and *φ*_*p*_ and *φ*_*s*_ are the derived prefix and suffix feature maps. Thus, once an FNN is trained to predict mAb-pMHC affinities, we can factorize it along any boundary we choose and obtain appropriate prefix and suffix embeddings. These embedding functions can then be used to guide the optimization of factorizable libraries.

### 2.3 Training data generation for a model of non-specific pMHC binding

Our campaign to train a model of non-specific pMHC binding was carried out by generating training data using three pMHC targets. The MHC molecule for all three targets is HLA–A*02:01, while the complexed peptides are Mage-A1 (KVLEYVIKV), Homeobox protein (NLQGSPVYV), and Tax Variant TM 10 (TLWG-WVKYV). We will refer to these targets as Mage, Homeobox, and Tax. Our initial phage display campaign was carried out in the context of a single framework library, FW-a. FW-a was used as the seed library for our experiments and was constructed as follows: a gene fragment encoding the germline framework combination IGHV3-23 and IGKV1-39 was synthesized in Fab format and cloned into a phagemid vector template. To generate training data CDR-H3s were diversified via trinucleotide synthesis technology, while the rest of the framework was held constant [10].

Our training campaign consisted of 14 experiments, each with a different round selection strategy using Mage, Homeobox, and Tax (Methods 5.1). All 14 experiments started with our FW-a seed library. The experimental selection rounds were performed as described in prior work [10]. As a final denoising step, sequences that had 5 or fewer observations in all 14 sequenced selection experiments were discarded.

Phage display CDR-H3 count data were then processed with our affinity inference procedure. For our prior, we selected normal distributions for log-initial concentrations with mean zero and variance 2. For log-affinity we chose normal distributions with mean zero and variance 1. The same seed library was used throughout this training campaign. However, instead of considering the initial concentrations to be the same random variable throughout all experiments, we instead split the experiments into 5 groups that each have a single initial concentration, which offers us additional flexibility to model how library concentrations might vary across experimental conditions. These 5 groups correspond to the structure of the phage display campaign, which is provided in Methods 5.1. To maintain some notion that these concentrations still arise from the same library, we set a covariance of 1 between each pair of initial concentrations if they are for a single sequence.

Using our inference procedure we computed non-specific mAb affinities for each CDR-H3 to train an FNN objective function to minimize for candidate CDR-H3 sequences. At this point sequences containing cysteine and asparagine were removed before training the FNN because of their poor representation in the dataset which causes their properties to be poorly inferred.

## 3 Results

### 3.1 Factorizable Neural Networks are a viable architecture for predicting non-specific affinities

Since FNNs are simple neural networks we wanted to ensure that they are capable of modeling the non-specific affinities of CDR-H3 sequences. To test this assumption we trained and evaluated an FNN and eight other baseline architectures including multilayer perceptrons with fully connected layers, convolutional resnets [7] and transformers [16] on their ability to predict log non-specific affinity, using the non-specific affinities we computed on CDR-H3 sequences as training data. Additional architectural details can be found in Methods 5.6.

Our optimization objective is to minimize the mean squared error loss between the predicted and inferred log-affinity values. To take advantage of the uncertainty estimates we have for our log-affinities, during training we sample minibatches with replacement, such that the probability of drawing a sample is proportional to the inverse of its variance. Our optimization objective is then equivalent to the likelihood over a heteroscedastic set of Gaussians. A minibatch consists of 512 samples, an epoch consists of 100 minibatches, and 200 epochs of gradient descent were performed using Adam [8] with default parameters.

We then compared the performance of FNNs and our baseline models on predicting log non-specific affinity and found that FNNs provide comparable performance to the baseline models. For validation, we created 100 different random training, test, and validation splits in ratios of 18 to 1 to 1. On each set of splits we trained all 9 architectures. Models were trained on the training set and tested on the test set after each epoch of training. The model obtaining the highest test performance was saved and scored on the validation set. The performance of the models are shown in Figure 1, where we can see that FNNs are comparably viable for the task of predicting nonspecific affinity as contemporary architectures, only slightly outperformed by transformer architectures, which are highly compute intensive and do not have the factorizable property of FNNs. We use weighted Pearson and Spearman correlations to evaluate our models, once again leveraging the uncertainty in our affinity estimates. Details on these metrics are provided in Methods 5.7.

**Fig. 1.**
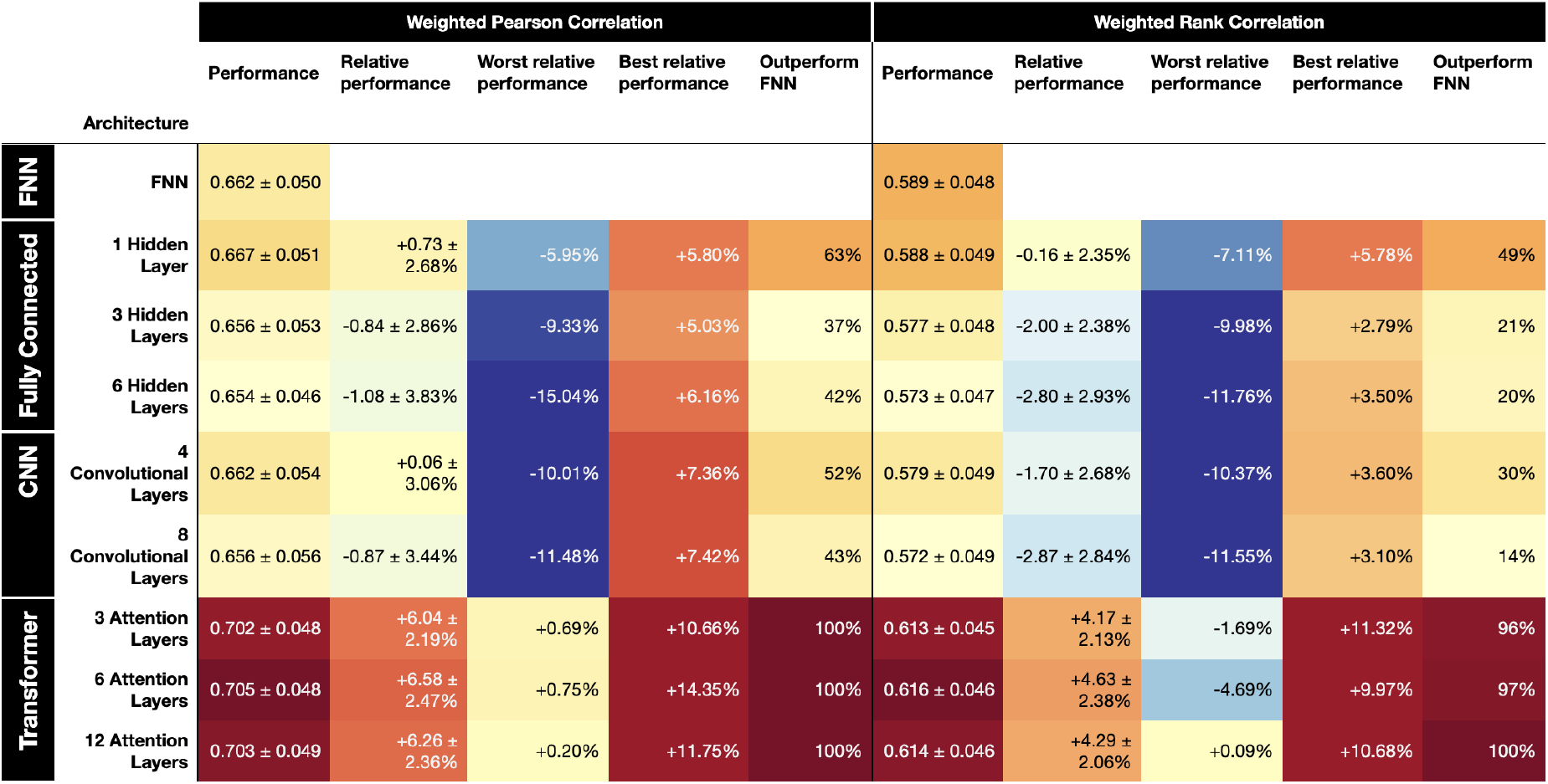
FNNs are competitive with contemporary deep learning architectures at predicting non-specific affinity. We evaluate the weighted Pearson and rank correlations between variationally inferred and predicted log-affinities. We assess the predictions made by FNNs, along with 8 other contemporary architectures: 3 multilayer preceptrons with fully connected layers (Fully Connected), 2 convolutional neural networks (CNN), and 3 transformers (Transformer). Two metrics are evaluated: Pearson correlation and rank correlation (also known as Spearman correlation). Within a given metric, the first column gives the raw performance metric with one standard deviation. The second column gives the relative performance achived by the architecture over the FNN, along with one standard deviation. The next two columns give the worst and best relative performances achieved out of the 100 trials, and the last column gives the percentage of trials in which the architecture outperforms the FNN. Out of these architectures, only the transformers achieve a level of performance that convincingly exceeds that of the FNN, and only the 12 layer transformer is able to beat the FNN in all 100 trials on both performance metrics.

We then constructed an FNN model to implement the our optimization objective to minimize log non-specific affinity. To construct our optimization objective, we trained an ensemble of FNNs. We first split our dataset into 10 roughly equal partitions by randomly assigning each sequence one of 10 numbers. This defines 10 pairs of training and test sets: a given partition is a test set, and the remaining sequences form the training set. For each training set and test set pair, we trained 10 FNNs on each split for 1000 epochs. The model attaining the best weighted Pearson correlation on the test set was selected from each partition, forming an ensemble with 10 members. The ensemble prediction is taken to be the mean of the prediction of its members. We then normalized the output such that it has zero mean and unit variance over the entire unpartitioned dataset.

### 3.2 Designing a library to minimize non-specific pMHC binding

Using our FNN model of log non-specific pMHC affinity we can now design a *de novo* CDR-H3 factorizable library that minimizes non-specific binding. Figure 2 depicts the design process for creating our baseline, enumerated, and factorizable libraries. Each library was given a design budget of 20,000 synthesized DNA molecules, and was designed using data from our experimental training data campaign. Each of the final libraries was tested in our evaluation campaign against the Flu GL9 and Mage-A1 pMHC targets as described in the next section.

**Fig. 2.**
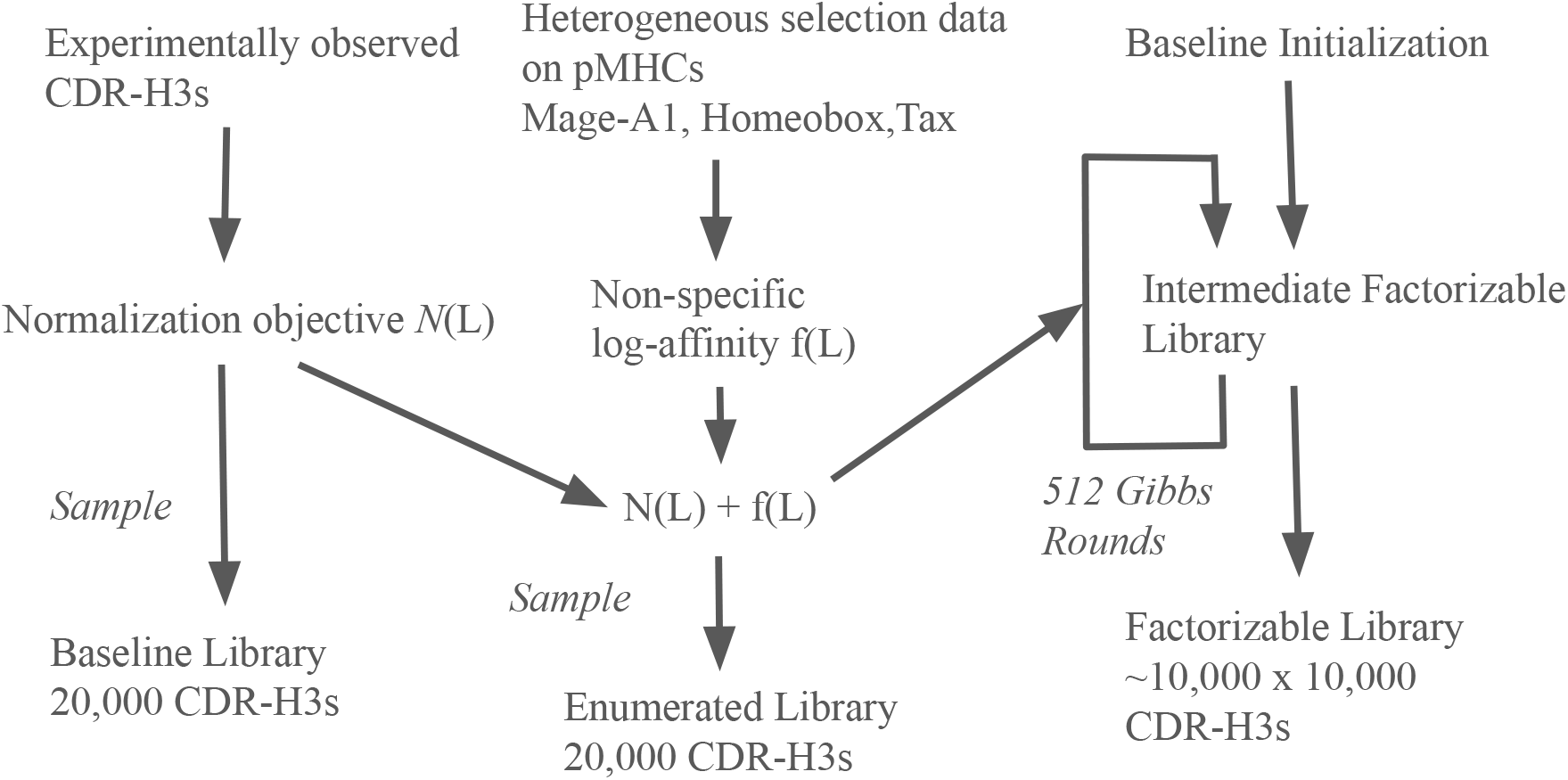
Design of the Baseline, Enumerated, and Factorizable libraries, each with a DNA synthesis budget of 20,000 sequences.

While it is possible to generate libraries with variable length CDR-H3 sequences, we chose to keep our sequences at a constant length of 16, the most frequently represented sequence length in our dataset. Furthermore, to keep our designs in a space where our models are reasonably confident, we remove from consideration certain residues at specific positions (Methods 5.8).

Direct optimization against surrogate functions described by deep learning models tends to result in adversarial examples that are meaningless outputs that the model scores highly [14]. To mitigate this, we introduce normalizing objectives to gravitate our library statistics toward more naturally observable distributions.

We model the naturally observable baseline distribution of CDR-H3 sequences with a Potts model [9], a generative model that is a generalization of the Ising model from statistical mechanics (Methods 5.9). To train the model we subjected the seed library FW-a to a single round of mock phage panning. This was repeated 5 times and the reads were combined (Methods 5.2). A further denoising step was taken where any CDR-H3 sequence with less than 5 observations in total were discarded. We then discard sequences containing cysteine and asparagine and sequences that are not of length 16, yielding in a total of 24890 unique sequences representative of a viable distribution of CDR-H3 sequences. We will refer to the uniform distribution over these unique sequences as the *baseline distribution* which we call *Q*. We first train our Potts baseline model on the observed baseline distribution *Q* and then construct a normalization objective 𝒩 (*L*) that we use to shift sampling distributions towards the distribution of viable sequences described by the Potts model (Methods 5.9).

Our full optimization objective, which we aim to minimize, evaluated on a library *L* is given by

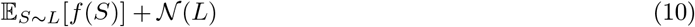

where *f* is the non-specific log-affinity as predicted by the FNN ensemble (larger is higher non-specificity). The objective N(*L*) increases as CDR-H3 sequences diverge from our trained observable baseline sequence model (Methods 5.9). The 𝔼 _*S*∼*L*_[*f* (*S*)] term encourages our designed library to have low non-specific log-affinity. Importantly, Potts models are factorizable as a product of a prefix embedding and a suffix embedding much like an FNN is, which allows us to evaluate 𝒩 (*L*) using the reverse kernel trick.

We design our factorizable library starting with an initial library that is evolved with simulated annealing. For our initial library, we drew sequences randomly from the *PWM* (*Q*) distribution, a distribution that is similar to the baseline distribution *Q* (Methods 5.9). Duplicates and sequences with residues in eliminated positions are discarded, and drawing continued until we obtained 10000 sequences. The sequences were split into prefixes of length 7 and suffixes of length 9. Suffixes were chosen to be longer due to the low variability in the last few residues. The initial library was then subjected to 512 rounds of Gibbs sampling, starting at an initial temperature of 100 for the first cycle and exponentially lowering it until the 512th cycle, where the temperature becomes 1*/*5000. Thus over the course of the optimization the draws become progressively more biased towards better scoring libraries by lowering the temperature parameter (Methods 5.10).

The resulting factorizable library consisted of 10000 prefix and 10000 suffix segments, which together could theoretically produce a library containing 10^8^ sequences. These segments were synthesized and joined via blunt end ligation. We sequenced the product, yielding 3432689 pairs of reads, of which 1739289 (50.67%) were successfully matched to library sequences. Within those sequences, 10000/10000 of the expected prefixes were observed, and 9993/10000 of the expected suffixes were observed.

To evaluate the relative performance of our factorizable library we designed two control libraries, where each control contained the same number of synthesized sequences as the factorizable library (20,000). The first control was an enumerated library that was designed with same objective as the facorizable library described by Equation 10, and thus tests the ability of the increased complexity of a factorizable library to produce additional candidates. The second control was a baseline library that was designed with only the normalization objective given in Equation 35. To produce libraries from these objectives, we first convert them into energy values defined sequence wise by thinking of sequences as libraries of size 1. We also multiply the objectives by the denominator of Equation 35. We then define a Boltzmann distribution over those energy values, from which we sample via Gibbs sampling. Each control library contained 20000 sampled sequences to match the number of sequences synthesized for the factorizable library.

### 3.3 A factorizable library produces more specific Flu-A02:01 binders than baseline controls

Given the designed baseline library, enumerated library, and factorizable library, we conducted a validation phage panning campaign against two peptide targets complexed with HLA-A02:01: Flu GL9 (amino acid sequence GILGFVFTL) and Mage-A1 (KVLEYVIKV). We will refer to the newly introduced Flu GL9 peptide as Flu. We expect that Flu binders in the enumerated library and the factorizable library will have lower Mage affinity, as a consequence of the anti non-specific objective used during library design. We further expect that the greater diversity of the factorizable library will enable us to observe a greater number of promising candidates. Data from Flu was not observed during construction of any of the three test libraries, and the objective is to see if the test libraries can produce more superior binders to Flu than Mage-A1. Data from Mage-A1 contributed to our non-specific log-affinity model.

The validation campaign evaluated the baseline, enumerated, and factorizble libraries to determine specific and non-specific mAb binding to Flu and Mage (Methods 5.1). Each experiment subjected the library under evaluation to at least two rounds of phage panning with Flu as the target, therefore we expect the resulting reads to have some bias towards Flu affinity. We converted read counts to affinity values (Section 2 and Methods 5.5). CDR-H3 affinities were not decomposed into separate specific and non-specific components, instead we extract raw Flu and Mage CDR-H3 log-affinities. Affinity inference was performed jointly for all three libraries to allow the inferred affinities to be comparable across different libraries.

We selected normal distributions for log-initial concentrations as our prior with mean zero and variance 2. For log-affinity we chose normal distributions with mean zero and variance 1. Each library was provided its own set of initial concentration variables. Unlike with the training campaign where a common seed library was shared throughout all experiments, no correlations were introduced to the priors here since different initial concentration variables correspond to the three different tested libraries.

After inferring Flu and Mage log-affinities, we assigned each sequence to its library of origin (baseline, enumerated, or factorizable library). The library designs contain no overlap, so each sequence was assigned unambiguously. Sequences that could not be matched to any library were discarded at this stage.

The 10 most significant sequences as scored by an adjusted Mahalanobis distance are given in Figure 4. To identify promising library candidates, we score each sequence based on its Flu and Mage log-affinity using an adjusted Mahalanobis distance which takes into account both how significantly a sequence’s Flu and Mage log-affinities deviate from the norm and how uncertain we are about those affinities. Sequences with high uncertainties are assigned lower significance (Methods 5.11). The distribution of log-affinities are provided as scatter plots in Figure 3, where the size of each point is proportional to its adjusted Mahalanobis distance.

**Fig. 3.**
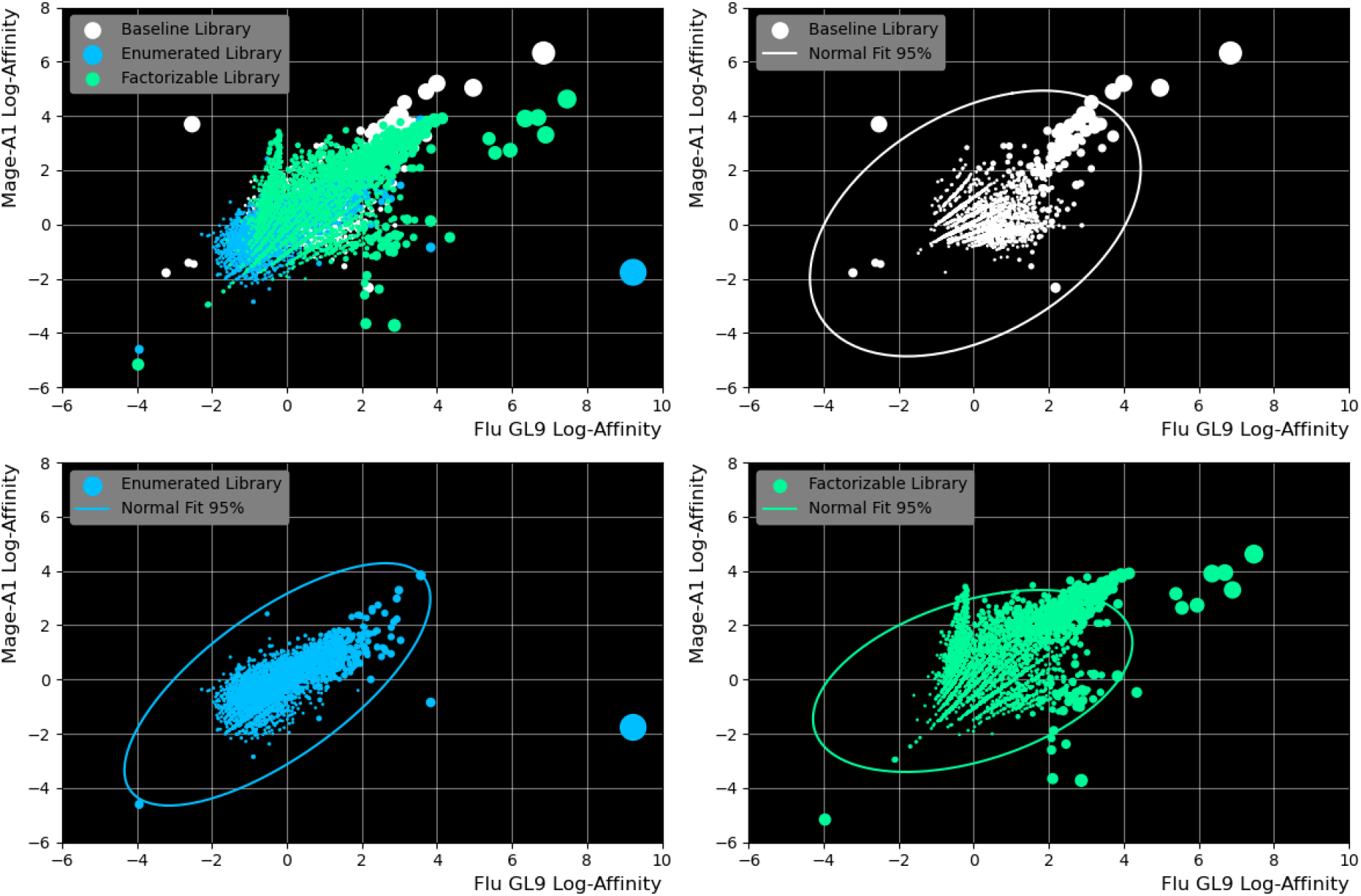
Scatter plot showing the Flu and Mage log-affinity distributions of the libraries. The Normal Fit 95% line indicates the isoprobability contour of a bivariate normal distribution that is fitted to the library distribution. The line is drawn so that 95% of the fitted distribution falls within the contour.

**Fig. 4.**
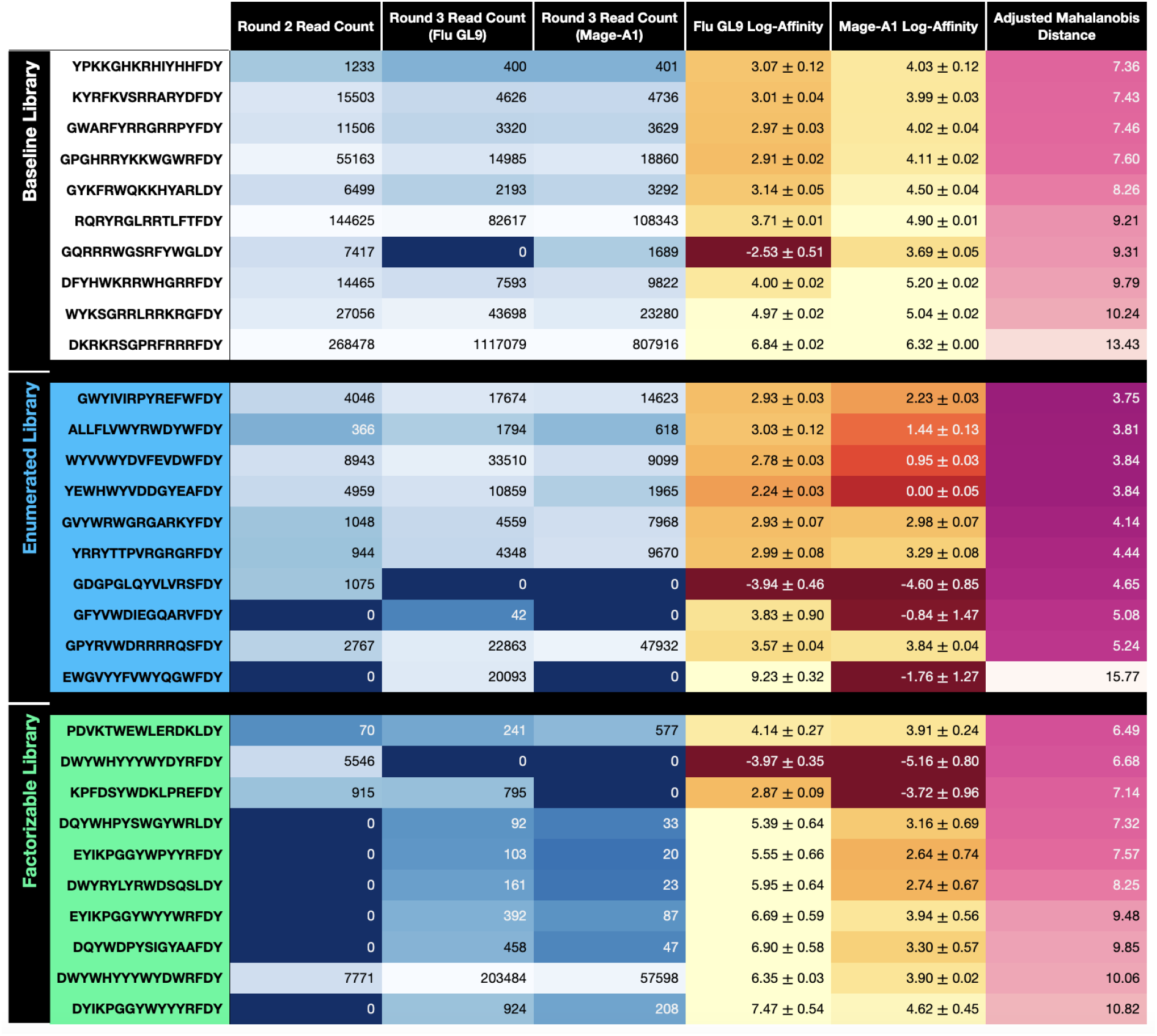
Most promising candidates for each library. Rows are sorted by adjusted Mahalanobis distance. One standard deviation is given for inferred log-affinity values.

We find the factorizable library contains a greater number of promising candidates (for example KPFDSY-WDKLPREFDY) than the control libraries with Flu log-affinities between 2-4. Thus, our hypothesis that a factorizable library produces more candidates than enumerated or random libraries with the same number of synthesized segments is supported. Observing the sequences with high Flu affinity (right side of the plot in Figure 3), we can see that the baseline library has higher Mage affinity, indicating that our non-specific design objective is successful in reducing the non-specificity of promising candidates. Referring to both Figure 3 and 4, we can see that the most promising candidate is EWGVYYFVWYQGWFDY in the enumerated library.

## 4 Discussion

Our primary contribution is the introduction of a practical method of designing high-complexity factorizable libraries that are shaped with a specified objective function. We used an objective function that penalized non-specific binding to peptide-MHC complexes, and for a fixed DNA synthesis budget we showed that factorizable libraries produced more specific peptide-MHC binding candidates than random or enumerated library controls. To produce an objective function that penalized non-specific binding we developed methods to extract non-specific components of affinity from heterogeneous high-throughput selection experiments. We showed how to use these data to design high-complexity factorizable libraries that are depleted of non-specific binders.

The design of factorizable libraries was enabled by the introduction of factorizable neural networks (FNNs) to represent a desired objective function in a manner that permits the efficient design of prefix and suffix segments. Sets of prefix and suffix segments that are randomly combined are designed according to an combinatorial objective that is described with an FNN. FNNs embed prefix and suffix sequences to enable the rapid recomputation of a combinatorial library prefix-suffix objective function when a single prefix or suffix segment is changed. The embedding thus admits an efficient method of sampling novel libraries according to a specified objective.

## 5 Appendix

### 5.1 Details on the training and validation phage display campaigns

We conducted two separate campaigns: a training campaign to obtain data to train a non-specific affinity objective function, and a validation campaign to evaluate the performance of our designed libraries. The read distributions obtained from the training campaign are summarized in Figure 5. Those obtained from the validation campaign are described in Figure 6.

**Fig. 5.**
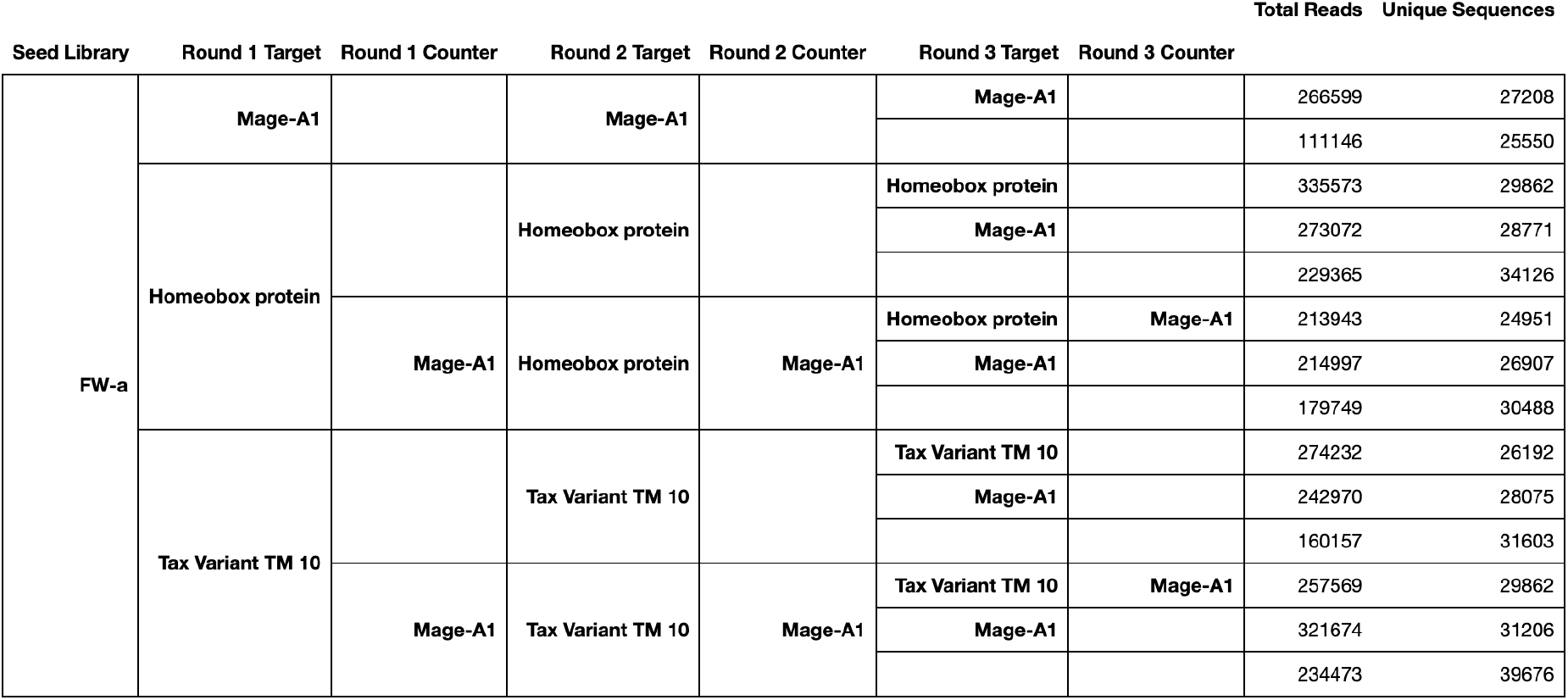
Phage panning experiments conducted in the training campaign. The training campaign consisted of fourteen different selection experiments corresponding to the fourteen table rows. A blank cell in Round 3 Target indicates no selection. The read counts given are those after removing sequences with less than 5 total counts across the entire campaign. For affinity inference purposes, all sequencing outputs that have the same round 1 and round 2 target and counterselection targets are assumed to have the same initial concentration value, giving a total of 5 initial concentration values.

**Fig. 6.**
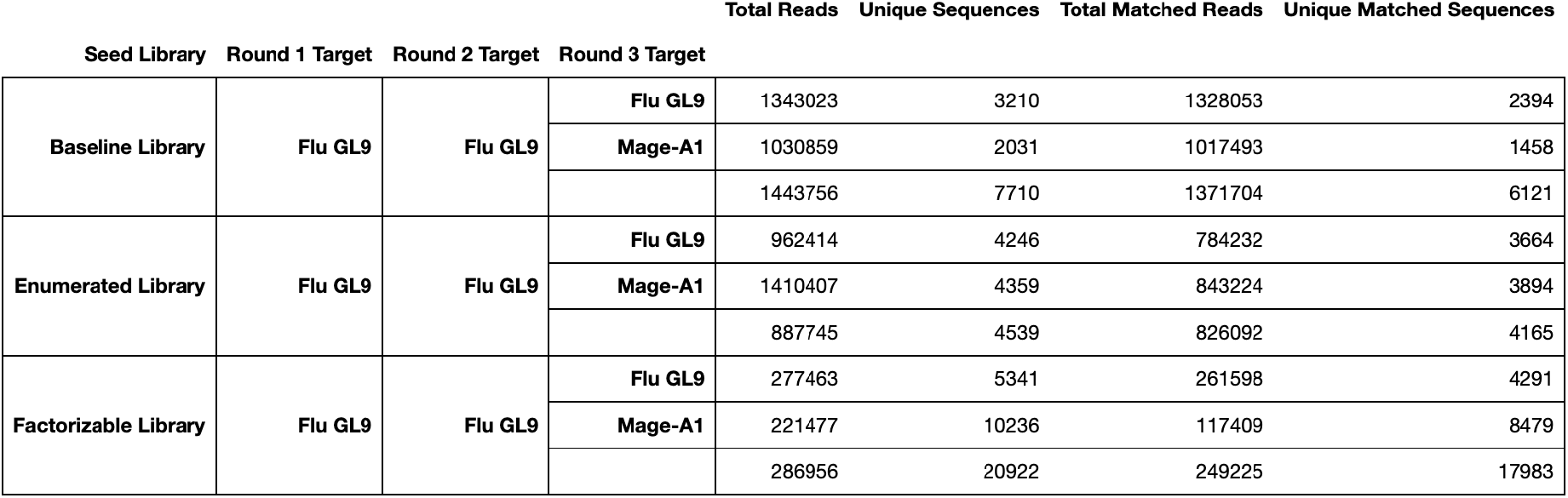
Phage panning experiments conducted in the validation campaign. The validation campaign used a total of nine experiments to evaluate three libraries for specific (Flu GL9) and non-specific (Mage-A1) binding. Instead of denoising by removing sequences with low observations, we match sequences to their seed libraries. For factorizable libraries, we checked to see if the prefix and suffix segments are both present. The resulting columns are given as “Total Matched Reads” and “Unique Matched Sequences”. For affinity inference purposes, all sequencing outputs that have the same seed library are assumed to have the same initial concentration value, giving a total of 3 initial concentration values.

### 5.2 Processing raw sequencing reads

High throughput sequencing is performed via MiSeq or HiSeq sequencers to assess the distribution of phages after a given number of rounds. We process sequencing data by first matching each read to the following regular expression:

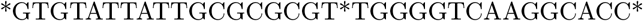

This represents the sequence flanking the variable CDR-H3 sequence in the framework of our seed library.

The wildcard in the center represents the variable CDR-H3 sequence itself.

We match the regular expression by first finding a sequence *s* that both matches the regular expression and has the minimal edit distance to the read. The part of *s* that matches the wildcard in the middle of the expression is defined as the extracted read, and the number of mismatches between *s* and the original read is defined as the number of errors in that read.

A graph is then constructed from unique reads, where an edge is formed between any pair of reads whose edit distance is 3 or less. Connected components are then extracted from the graph. We define a read as valid if its length is a multiple of 3 and if it contains no uncalled bases. If more than half the reads represented in a connected component are identical and valid, then the rest of the sequences in the component are corrected to the majority sequence. If this condition is not met, then no correction is made. All invalid sequences are then discarded. Sequences are then translated into their CDR-H3 amino acid sequences, and sequences containing any stop codons are discarded. Furthermore, we discard every unique CDR-H3 sequence where the average number of read errors across all reads with that sequence exceeds 4, and any CDR-H3 sequence that is shorter than 10 amino acids or longer than 20 amino acids in length. After processing we obtain a set of CDR-H3 sequences and the number of times they occur in the sequencing data (i.e. their read count), which represents the mAb distribution.

### 5.3 Learning affinities from phage panning read counts

Our first task is to estimate individual antibody affinities from phage display read counts. This objective is complicated by the diverse and unknown concentrations of antibodies in a candidate library. Thus observed antibody read counts cannot be directly interpreted as affinities.

Let [*P*_*s*_] be the concentration of phages displaying a mAb *s* at the end of the round. Let [*T*] be the concentration of free target at the end of the round. Let [*TP*_*s*_] be the concentration of phages displaying *s* that are complexed with the target. Assuming the concentration equilibriates during the wash we can express the affinity between *s* and the target (denoted *K*_*T,s*_) as the following:

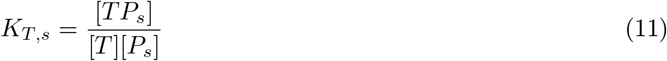

The concentration of phages displaying *s* at the beginning of the round is the sum of complexed and free phages at the end of the round. By rearranging this relation and substituting [*P*_*s*_], we find that the concentration at the start of a round ([*P*_*s*_] + [*TP*_*s*_]) and the concentration at the end of a round ([*TP*_*s*_]) is related by:

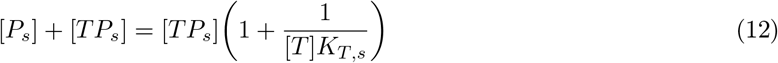

To model counterselection, recall that during counterselection an additional target is added which is not fixed, and is washed away after the solution equilibriates. To model this, let [*N*] denote the concentration of the counterpanning target at the end of the phage display round, and let [*NP*_*s*_] denote the concentration of phages that display *s* that are complexed with the counterpanning target. Again, assuming the wash equilibriates, the affinity between *s* and the counterselection target, which we denote *K*_*N,s*_, can be expressed as:

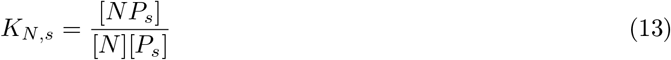

The concentration of phages displaying *s* at the beginning of the round is then the sum of complexed and free phages at the end of the round. By combining Equation 11 and Equation 13, we can solve for [*NP*_*s*_] as a function of [*TP*_*s*_]. Substituting [*NP*_*s*_], we can derive that the concentration at the start of a round ([*P*_*s*_] + [*TP*_*s*_] + [*NP*_*s*_]) and the concentration at the end of a round ([*TP*_*s*_]) is related by:

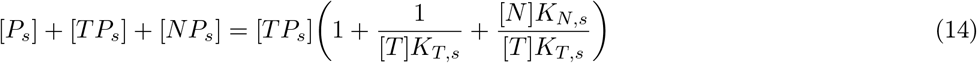

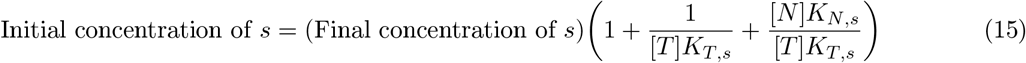

To complete our model, we make two simplifying assumptions:

1. We assume that the final concentration of a mAb at the end of one round is directly proportional to the initial concentration of a mAb at the start of the next round.
2. We assume that the read count of a particular mAb in the sequenced data follows a Poisson distribution whose parameter is directly proportional to the final concentration of that mAb in the round of phage display directly preceding the sequencing.

The first assumption allows us to extend the relation we derived relating the initial and final concentrations of a single round to a relation between the concentrations at any arbitrary round. The second assumption allows us to connect the read counts we obtain to these parameters. With these assumptions, the read count of *s* after *n* rounds of phage panning is Poisson distributed with parameter:

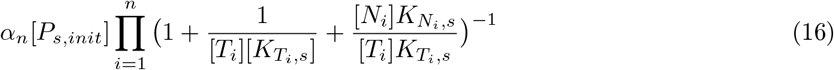

Where *α*_*n*_ is some scaling parameter that relates read count to concentration, [*P*_*s,init*_] is the initial concentration of phages displaying *s, T*_*i*_ is the panning target on the *i*th panning round, *N*_*i*_ is the panning target on the *i*th panning round. Given a set of observed read counts, we can then compute the likelihood of those observations using this relation, which we can optimize over.

We use variational inference to obtain approximate posterior distributions over log-affinity and log-initial concentration for each antibody CDR-H3. This permits us to quantify both an affinity value and an uncertainty around that value. This is important because although some mAbs can have very large round-over-round read counts, indicating high affinity, those increases may be supported by relatively few reads, indicating high uncertainty. Variational inference is performed by finding distributions *Q* over log-affinity and log-initial concentration from parametrized multivariate normal distributions to maximize the following Evidence Lower Bound (ELBO):

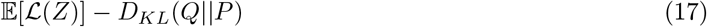

We maximize the ELBO over *Q*, where *P* is a prior distribution over the parameters and 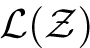 is the data likelihood of the observed read counts. ℒ (*Z*) is calculated by replacing *λ*_*s,n*_ with Equation 16 in the following formula for log Poisson distributions:

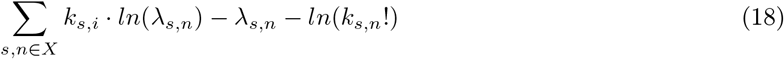

where *k*_*s,n*_ is the read count of *s* after *n* rounds of phage panning. Note that for optimization purposes the final term is irrelevant and can be discarded.

As mentioned, we enforce *Q* such that under it the log-affinity values *K* and log-initial concentrations are normally distributed, and that the remaining parameters are distributed as point distributions (Dirac Delta). We permit correlations between log-affinities and log-initial concentrations for a single mAb *s*, but not across different mAbs for computational efficiency. We use normal distributions as priors *P* for log-affinity and log-initial concentrations, and flat distributions for the remaining parameters.

### 5.4 Decomposing affinities into specific and non-specific components

We can decompose mAb affinity into specific and non-specific components by simultaneously solving for these compontents in the context of multiple experiments that have been designed to resolve specificity using different selection strategies. A key advantage of our model based approach to attaining affinity values is that it allows us to take phage display data with heterogeneous targets and counterselection targets and infer affinity using a single consistent model. This integration of heterogeneous target affinity information further allows us to decompose our affinity values into specific and non-specific components at the inference stage.

In this work, we are primarily concerned with mAbs that bind to peptide-MHC (pMHC) targets. Specifically, we would like mAbs that are specific to the peptide part of the complex, instead of binding indiscriminantly to arbitrary pMHCs. We expect that the non-specific binding we observe occurs through binding poses that ignore the peptide. If an mAb does not bind in part to the peptide in the MHC groove then the binding interaction is independent of the identity of the peptide that is complexed with the MHC. In the presence of non-specific binding we assume that we have the following possible reactions:

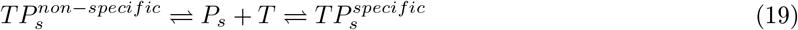

Where 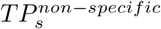 is the complex formed through non-specific interactions and 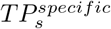 is the complex formed through specific interactions. Assuming that these are the only reactions that can occur (i.e. we cannot directly shift from one complex to another in a single step), we have two affinities at equilibrium:

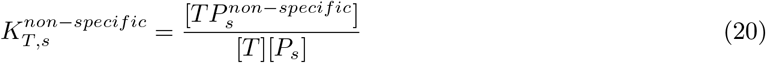

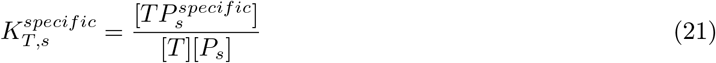

Rearranging this then yields:

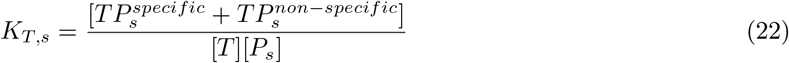

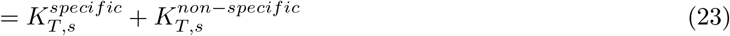

Where *K*_*T,s*_ is the overall affinity, and 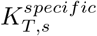and 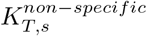 are the specific and non-specific affinities respectively. Therefore, we can simply substitute *K*_*T,s*_ with 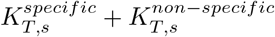 in Equation 16 to account for the different components of the affinities, impose a prior over both, and carry out inference as before to attain an isolated non-specific affinity.

### 5.5 Technical notes on optimizing the Evidence Lower Bound

To calculate the ELBO itself requires taking an expectation which is non-trivial to compute. Here, we estimate the ELBO via sampling. 2048 samples are drawn from the parametrized distributions to estimate the ELBO. Only sequences with non-zero read counts were used in this calculation with the exception of cases where two sequencing datasets shared a round of phage display, but one dataset contained a sequence that was not observed in the other dataset. In this case that sequence was assigned a read count of zero for the other dataset, and was included in the calculation of the ELBO. The parametric distributions are then optimized by backpropagating the gradient from the ELBO to the distribution parameters and performing gradient descent. Adam [8] was used for descent with default parameters with the exception of learning rate, which was set to 0.1, and 10000 iterations were performed. The optimization was repeated 5 times, and the parameters achieving the best ELBO across the 5 repeated runs and across all 10000 iterations of the runs were taken as the final parameters.

### 5.6 Model architecture details

Including FNNs, a total of 9 architectures were trained:

– Factorizable Neural Network - a three layer perceptron with exponential activation and 128 hidden units.
– 1 Hidden Layer - a three layer perceptron with ReLU activation and 128 hidden units.
– 3 Hidden Layers - a multiplayer perceptron with ReLU activation and three hidden layers, which have 512, 128, and 32 hidden units in order of a forward pass.
– 6 Hidden Layers - a multiplayer perceptron with ReLU activation and six hidden layers, which have 512, 512, 512, 256, 128, and 32 hidden units in order of a forward pass.
– 4 Convolutional Layers - a 1D convolutional neural network with residual connections between convolutions. Kernels are of size 3. The network contains a total of 4 layers, with downsampling by a factor of two after each layer achieved through average pooling.
– 8 Convolutional Layers - a 1D convolutional neural network with residual connections between convolutions. Kernels are of size 3. The network contains a total of 8 layers, with downsampling by a factor of two after every two layers achieved through average pooling.
– 3 Attention Layers - A transformer with 3 attention layers. Each attention layer has 8 heads. Keys, queries, and values are 512 dimensional. A fully connected layer then maps the output to a vector of 128 dimensions, which is then passed through a ReLU layer and then linearly transformed into the output.
– 6 Attention Layers - Same as 3 Attention Layers, but with 6 attention layers.
– 12 Attention Layers - Same as 3 Attention Layers, but with 12 attention layers.

These could broadly be categorized into 4 categories. The first model in the above list is the FNN, which is in its own category. The next three are multilayer perceptrons with fully connected layers. The next two are convolutional neural networks. The last three are transformers.

Due to the idiosyncracies associated with the architectures, the CDR-H3 sequences were encoded slightly differently for each category. The following is shared between encodings: for each amino acid (with the exception of cysteine and asparagine due to their exclusion from the dataset) we assigned a one-hot encoded vector. A padding letter was also introduced, and was also assigned a one-hot vector. The one-hot vectors were therefore 19 dimensional. Each sequence was padded to be length 20.

– For FNNs, padding was applied in the middle. In other words, or a length *L* sequence we took the first ⌊*L/*2⌋ letters, concatenated 20 − *L* padding characters, and then concatenated the remaining ⌈*L/*2⌉ characters. An additional character was appended to the end to denote the length of the sequence. Since we only consider sequence lengths between 10 and 20 inclusive, only 11 different lengths need to be represented, so we simply use the one-hot representations we already have for the amino acids^4^.
– For multilayer perceptrons with fully connected layers, we similarly apply padding in the middle of the sequence like for FNNs. Instead of denoting lengths with an additional character, we instead add an additional channel, giving us a total of 20 channels (19 that account for the one-hot vector and 1 that accounts for the length). The additional channel contains the same value at each entry, which we set to (1 + *e*^(10−*L*)*/*2^)^−1^.
– For convolutional neural networks, we pad out the end of the sequence. This is because having a gap in the middle would harm the models ability to share information between the part of the sequence before the gap and the part of the sequence after the gap. We then add an additional channel that encodes length information identically to how we do for multilayer perceptrons.
– For transformers, we encode sequences in nearly the same manner as we do for multilayer perceptrons. However, we add an additional 20 channels that encode position information, since transformers are otherwise largely position agnostic. 10 of the 20 channels contain the value *sin*(10000^−(2∗*i*)*/*20^*x*) where *x* is the position ranging from 0 to 19 inclusive, while *i* are the indices for the channels ranging from 0 to 9 inclusive. The other 10 channels contain the value *cos*(10000^−(2∗*i*)*/*20^*x*) where *x* is the position ranging from 0 to 19 inclusive, while *i* are the indices for the channels ranging from 0 to 9 inclusive.

### 5.7 Validation metrics for assessing deep learning models over datasets with variable uncertainty

For our testing/validation metric, we use an extension of Pearson and rank correlations that extends to weighted datasets, where weights are inverse variances. This way, a datapoint that we have high confidence in can be more important than a large amount of datapoints that we are not confident about.

We can extend Pearson correlation to weighted datasets in the following way: we can view a vector of datapoints as an empirical distribution over the elements in the vector. If the two distributions are denoted *X* and *Y*, Pearson correlation can be defined by:

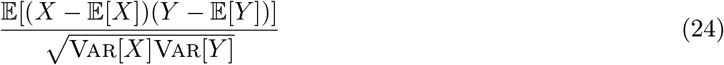

This definition then extends without modification to weighted datasets, which can be viewed as a weighted distribution.

We can take the same view for rank correlation. Given *X* and *Y*, we define *Φ*_*X*_ and *Φ*_*Y*_ as the cumulative distribution functions for the marginal distributions of *X* and *Y* respectively. Then rank correlation can be defined as:

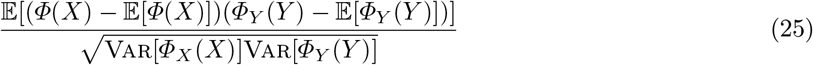

### 5.8 Eliminating position specific residues from consideration

We eliminated methionine from all positions due to its poor support in our training data. We further eliminated all residues from positions 14, 15, and 16, with the exception of tyrosine at position 16, aspartate at position 15, and phenylalanine and leucine at position 14, also due to poor support.

### 5.9 Construction of a Potts model to match the baseline distribution

A Potts model model defines a distribution by assigning an energy value to each sequence. The probability of observing that sequence is then proportional to the exponential of the negative of its energy. The energy can be scaled by a temperature parameter that further controls the shape of the distribution. The energy of a sequence under a Potts model is a sum of pairwise interaction energies:

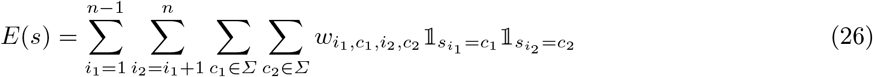

Where *s* is the input sequence of length *n, Σ* is the alphabet the letters in the sequence are drawn from, and the indicators equal 1 if the character at a given position in the sequence equals some given character. The model is parametrized by the interaction energies *w*.

Sampling from this model can be accomplished via Monte Carlo methods like Gibbs sampling, where we start from a sequence drawn uniformly at random. Then each character is resampled in sequence from the marginal distribution. The process is then iterated to converge to the stationary distribution. To fit a Potts model, we maximize the log likelihood of the observed sequences:

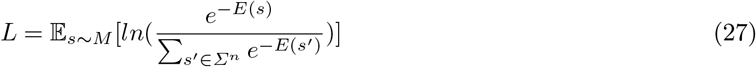

Where *M* is the distribution we wish to match, and *Σ*^*n*^ is the set of all sequences of length *n*. To maximize this quantity, we note that the first and second derivatives are:

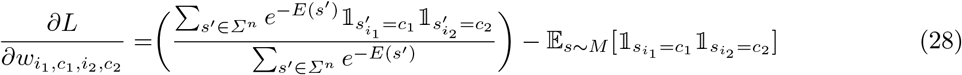

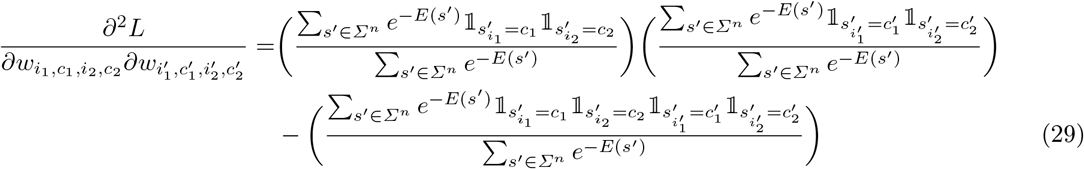

The terms in the large brackets above are probabilities that sequences drawn from the Potts model contain characters at given positions. Let *P* (*i*_1_ = *c*_1_, *i*_2_ = *c*_2_, …, *i*_*n*_ = *c*_*n*_) be shorthand for the probability that a sequence drawn from the Potts model contains *c*_1_ at position *i*_1_, *c*_2_ at position *i*_2_, and so forth. We can then rewrite the derivatives as:

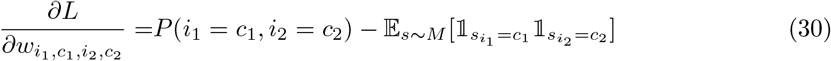

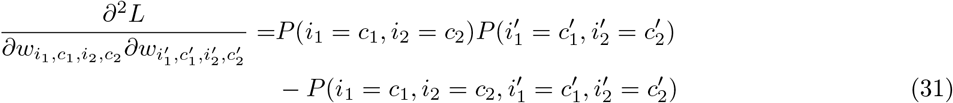

We would expect that as more conditions need to be satisfied, the number of sequences satisfying that property drops exponentially. Therefore, the terms dominating the second derivatives are:

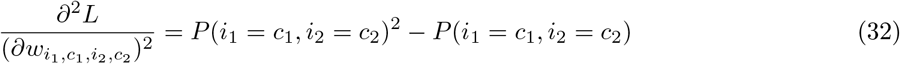

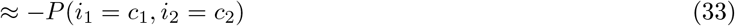

Because only two conditions need to be satisfied in this term. Taking this approximation and assuming that all other second derivatives are negligible, we can solve via Newton’s method: given a set of parameters *w*, we perform the following update:

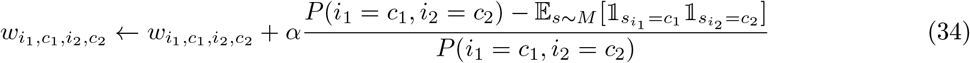

where *α* is the step size. For our optimization, we set *α* to 0.005. Calculating the updates also require us to calculate the Potts model probabilities of observing sequences with residues at given positions. We approximate these probabilities via 1048576 samples each obtained from running 100 rounds of Gibbs sampling on a uniformly randomly drawn seed.

This methodology was used to match a Potts model to the baseline distribution. Sampling from the final optimized Potts model yields pairwise frequencies that differ from pairwise frequencies observed in baseline distribution by no more than 0.001.

Let 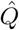 be the distribution described by the trained Potts model. Given any distribution *P* over a probability space of constant length sequences, let *PWM* (*P*) denote the distribution that is closest to *P* subject to the constraint that the symbol at each position along the sequence is independent of the others. Then given a library *L*, the normalizing objective 𝒩 we aim to minimize is:

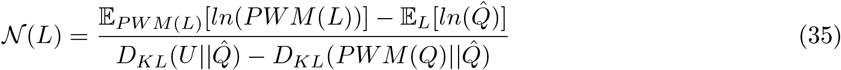

Where *U* is a uniform distribution over all sequences that are considered for library inclusion (see Section 5.8). Note that here we are viewing a library *L* as a probability distribution that is uniform over its members. If we remove the *PWM* operator, the numerator becomes the KL-divergence between the library distribution *L* and the Potts model distribution 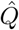. Therefore, applying this objective ought to result in a library distribution to similar to 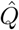. The denominator is for normalization purposes, so that the objective value moves by roughly a single unit when we move from a uniform distribution to one that resembles 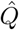.

### 5.10 Gibbs Sampling

Gibbs sampling a factorizable library is performed residue by residue in the following way: a specific residue from a specific sequence in the prefix and suffix library is chosen. All possible valid mutations of that residue are enumerated, provided that the mutation does not introduce a duplicate sequence into the library. The values of the new factorizable libraries given those mutations are computed via the reverse kernel trick. A Boltzmann distribution (i.e. a softmax over a set of negative energy values divided by a temperature parameter) is then defined over those mutations, where the energy value is the negative score of the factorizable library resulting from the mutation. A mutation (including no mutation) is sampled according to that distribution. We then proceed to the next residue and repeat the sampling process. Residues are iterated in columns: first, all first residues in prefixes are updated. Next, all final residues in suffixes are updated. Then all second residues in prefixes are updated, and so on. This continues until every single residue has been updated, at which point the cycle repeats with an updated temperature.

### 5.11 Adjusted Mahalanobis distance

The Mahalanobis distance can be used to detect outliers when working in a multidimensional setting. Given an empirical distribution of samples in ℝ ^*n*^, we first estimate its distribution by calculating the mean *mu* and covariance *Σ* of the samples, which defines a multivariate normal distribution.

The standard Mahalanobis distance of a sample *x*, denoted here as *D*(*x*), is then given as:

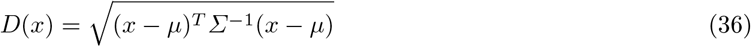

Which gives the amount by which a sample deviates from a typical sample. In our case, our samples come attached with their own covariances that quantify our uncertainty about those points. High uncertainty points ought to be given less significance, so we adjust the Mahalanobis distance for a sample *x* with covariance *Σ*_*x*_ to:

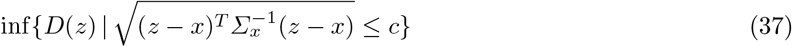

Where *c* ≈ 2.45 is chosen such that the measure of the set that the infimum operator is taken over is 0.95, approximately analogous to 2 standard deviations in the one dimensional case.

Intuitively, what this achieves is to find a realization of the sample we are uncertain about that is reasonable and provides the lowest possible Mahalanobis distance, thus diminishing the significance of samples we have high uncertainty about.

## Footnotes

4 This representation can then be flattened to adhere to the definition of an FNN given in Equation 4, although our implementation of an FNN works with the matrix input without flattening.

